# A Structural Landscape of Fungal Allergens with Epitope-Level Resolution Reveals Cross-Kingdom Structural Similarity

**DOI:** 10.64898/2026.01.24.701546

**Authors:** KyuTark Kim

## Abstract

The immunological properties of allergens are ultimately governed by three-dimensional protein structure, particularly the spatial organization of IgE-binding epitopes. Despite the need to capture global structural relationships and patterns of allergen diversity, most allergen classification and prediction strategies still rely predominantly on sequence-based approaches. Here, we present a structure-based framework that represents fungal allergens as elements embedded within a continuous structural manifold. Using AlphaFold-predicted full-length structures of 150 previously reported fungal allergens, we constructed a global structural distance space defined by TM-score-derived similarities. This manifold revealed a heterogeneous yet continuous landscape in which dense structural neighborhoods correspond to established allergen families, while more diffuse regions reflect gradual structural transitions. To directly link global protein architecture with immunologically relevant features, we extended this framework to epitope-level structural representations derived from predicted antibody-binding regions. Epitope-restricted structures preserved the relative organization of major allergen clusters, demonstrating that IgE-relevant features are embedded within conserved structural scaffolds. At the same time, epitope-level analysis demonstrated that high structural similarity can be observed at the epitope level even when global protein folds differ, highlighting the contribution of local structural features to immunologically relevant properties. We further applied a nearest-neighbor-based manifold inclusion analysis to screen nearly 20,000 fungal protein structures, identifying numerous allergen-related proteins occupying the same structural neighborhoods as known allergens, thereby extending allergen-associated architectures beyond current databases. Finally, structural screening of non-fungal allergens revealed cross-kingdom similarities to fungal allergens, suggesting convergent allergenic architectures across distant taxa. Together, this study establishes a unified structural manifold framework that integrates full-length protein and epitope-level information, providing a structure-based perspective on allergen diversity as a continuous space and on cross-reactivity.

## 1. Introduction

The immunological properties of allergens are fundamentally determined by the three-dimensional organization of the protein, and in particular by the structural features of IgE-binding epitopes^1^. Although allergic sensitization and immune responses ultimately arise from structural recognition at the epitope level, most existing allergen classification and prediction strategies have relied primarily on sequence similarity or sequence-derived domain annotation frameworks^2^. While effective for identifying closely related allergens, these approaches are not designed to explicitly capture structural and functional relationships among more distantly related proteins, nor to provide a global, integrative understanding of allergen diversity.

Recent advances in deep learning-based protein structure prediction, exemplified by AlphaFold and related models, have enabled reliable generation of three-dimensional protein structures at large scale^3^. This technological shift provides an opportunity to reinterpret allergens not as discrete sequence families, but as elements embedded within a continuous structural space^4^. Structural similarity quantified independently of sequence identity more directly reflects the molecular reality of protein-protein interactions involved in immune responses, including antibody-antigen recognition, thereby offering a more biologically grounded framework for allergen analysis^5^.

Fungal allergens represent a clinically important class of allergens characterized by diverse sources of exposure, including food products and airborne fungal particles across environmental and indoor settings, and are implicated in a wide range of respiratory, food, and occupational allergic diseases^6^. This diversity of exposure routes and clinical manifestations suggests substantial heterogeneity among fungal allergens^7,8^. However, despite their clinical relevance and extensive investigation in specific contexts, a unified structure-based perspective that integrates their global diversity and interrelationships remains largely unexplored.

Here, we systematically analyzed all reported fungal allergens, including those with experimental validation, by constructing a pairwise structural distance matrix based on full-length allergen protein structures, thereby defining a global structural manifold of fungal allergens. This manifold-based representation enables allergen relationships to be interpreted as continuous structural neighborhoods rather than discrete categories. To directly link each protein’s global structural organization with immunologically relevant features, we extended this framework to epitope-level representations derived from predicted IgE-binding regions, and examined how density-based clustering patterns observed at the full-length level are preserved or reorganized at the epitope level.

Within this unified structural framework, we further assessed whether additional fungal proteins occupy structural neighborhoods proximal to known fungal allergens, enabling the identification of candidate allergen-related proteins beyond current databases. Finally, we examined structural similarity between fungal allergens and allergens reported from other biological kingdoms to explore the extent of cross-kingdom structural convergence.

Together, this study establishes a structure-based manifold framework grounded in full-length allergen protein structures and extends epitope-level structural analyses based on IgE-binding sites predicted from these structures. Through this framework, allergen diversity, epitope conservation, and cross-reactivity can be interpreted within a continuous structural space.

## 2. Methods

### 2.1. Datasets

Experimentally validated fungal allergens were collected from the WHO/IUIS Allergen Nomenclature database, COMPARE, and SDAP 2.0^1,9,10^. After removing redundant entries, a total of 150 fungal allergen proteins derived from 42 fungal species were retained, all of which had accessible protein sequences via UniProt or GenBank identifiers.

To identify potential novel fungal allergens, all fungal protein structures deposited in the Protein Data Bank were retrieved based on taxonomic annotation corresponding to the fungal kingdom (NCBI taxonomy ID: 4751). Protein structures were split at the PDB entity level, yielding 19,999 fungal protein entities derived from 3,502 PDB entries.

For cross-kingdom allergenicity analysis, 1,289 experimentally validated non-fungal allergens were obtained from the WHO/IUIS Allergen Nomenclature database^9^. These allergens originated from 295 species spanning the Animalia, Bacteria, and Plantae kingdoms, and corresponding protein sequences were retrieved via UniProt or GenBank identifiers for downstream analysis.

### 2.2. Protein Structure Modeling

Protein sequences were used for downstream structural modeling. Three-dimensional protein structures were predicted using AlphaFold implemented via ColabFold (v1.5.5) in batch mode with default parameters^3,11^. For structure prediction, multiple sequence alignments were generated using the default ColabFold MSA pipeline. Input FASTA headers were standardized to unique sequence identifiers based on the original UniProt or GenBank accession IDs, ensuring consistent mapping between sequences and predicted structures. For each sequence, a single representative structure was selected by prioritizing the top-ranked prediction (rank_001) for downstream analyses. Model quality was assessed using pLDDT scores (mean: 85.82), with 88.0% of structures showing high confidence (>70). Lower-scoring models were retained to capture intrinsically disordered regions characteristic of certain allergens (Supplementary Fig. 1).

### 2.3. IgE-Relevant Epitope Definition and Structural Representation

IgE-binding epitope regions of fungal allergens were predicted using BepiPred 3.0 and DiscoTope 3.0. For BepiPred, per-residue linear B-cell epitope scores were computed using default settings, and residues with scores greater than or equal to the recommended threshold (0.1512) were selected^12,13^. In parallel, structure-based conformational epitope prediction was performed using DiscoTope 3.0 on AlphaFold-derived structures, and residues classified as epitope-positive according to the calibrated DiscoTope criteria were retained.

Epitope residues identified by either method were combined, and the corresponding atomic coordinates were extracted from the full-length allergen structures. These residues were used to generate epitope-restricted structural representations for each allergen. The resulting epitope-restricted structures were used exclusively for epitope-level structural comparison and interpretation, while full-length structures were retained for primary manifold construction.

### 2.4. Construction of the Allergen Structural Manifold and Clustering

Pairwise structural similarity among the 150 fungal allergens was quantified using TM-align (version 20240303)^14^. For each protein pair, the two TM-scores reported by TM-align were averaged to obtain a symmetric similarity measure. This similarity was then converted into a distance metric defined as *D* = 1 − *TM*, yielding a 150 × 150 symmetric distance matrix that served as the primary fungal allergen structural manifold. Hereafter, the term “structural manifold” is used to denote a distance-defined relational space constructed from pairwise TM-align similarities.

Clustering of the structural manifold was performed using HDBSCAN with a precomputed distance metric^15^. To visualize the global organization of allergen structures, two-dimensional embeddings were generated using t-SNE applied to the precomputed distance matrix^16^. Clusters of full-length allergen structures were labeled alphabetically in descending order of cluster size, while unclustered proteins were retained as noise.

As an additional biological analysis, epitope-restricted structural representations were used to evaluate epitope conservation within structurally defined allergen families. Pairwise similarity computation and distance definition for the epitope-restricted structures followed the same TM-align-based procedure (*D* = 1 − *TM*). Clustering and visualization were performed as for the full-length allergens to enable direct comparison between global structural similarity and epitope-level structural organization. Epitope clusters were labeled numerically such that epitope cluster indices corresponded to the alphabetical ordering of their originating full-length protein clusters.

### 2.5. Manifold Inclusion Analysis and Candidate Identification

To assess whether external proteins reside within the known fungal allergen structural manifold, a density-based inclusion criterion using nearest neighbors was defined using the reference allergen.

Using the reference structural distance space defined above, a nearest-neighbor density-based inclusion criterion was applied. Let *r*_1_ denote the nearest reference allergen to a query structure *q*. For each reference allergen *r*, a local neighborhood radius *τ*(*r*) was defined as the distance to its k-th nearest reference neighbor within the reference distance matrix (k = 3; self-distances excluded). A query was classified as residing within the manifold if its distance to the nearest reference allergen satisfied *D*(*q, r*_1_) ≤ *τ*(*r*_1_); otherwise, it was classified as outside the manifold.

This inclusion framework was applied independently to both fungal protein candidates and non-fungal allergen candidates. In both cases, candidate sequences were first identified by MMseqs2 sequence-based pre-filtering against the fungal allergen set using minimum coverage and sequence identity thresholds of 0.1^17^. All candidates were then evaluated using the same nearest-neighbor density criterion described above, enabling consistent assessment relative to reference fungal allergens.

## 3. Results

### 3.1. Fungal allergens occupy a continuous yet heterogeneous structural landscape

A global structure-based space of fungal allergens was constructed using full-length structures of 150 experimentally validated fungal allergens (Fig. 2A). Pairwise TM-score-derived distances among all allergens (11,175 pairs) revealed a broadly distributed structural landscape, with a mean distance of 0.73 and a median distance of 0.75. While the maximum observed distance reached 0.9148, a subset of allergen pairs exhibited very close structural similarity. The overall distance distribution showed moderate dispersion (SD = 0.11), indicating that fungal allergens occupy a continuous yet heterogeneous structural space.

**Figure 1.**
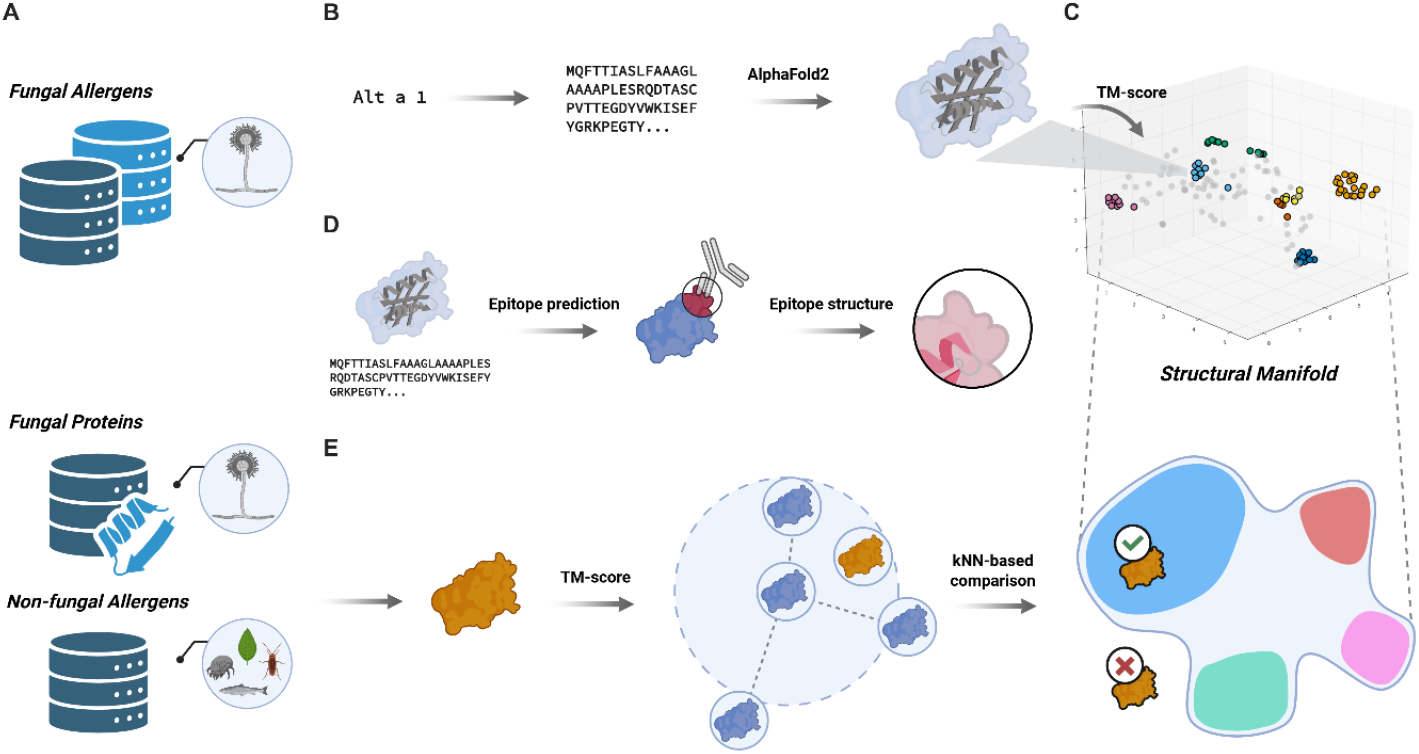
Overview of the structure-based framework for fungal allergen analysis. **(A)** Curated datasets of fungal allergens, fungal proteins, and non-fungal allergens used in this study. **(B)** Representative fungal allergen sequences were used for protein structure prediction using AlphaFold2. **(C)** Predicted full-length protein structures were embedded into a global structural manifold based on TM-score-derived structural similarity. **(D)** IgE-binding epitopes were predicted from full-length structures and used to generate epitope-level structural representations. **(E)** Query proteins were evaluated by kNN-based comparison within the structural manifold to determine manifold inclusion. Created with BioRender.com.

**Figure 2.**
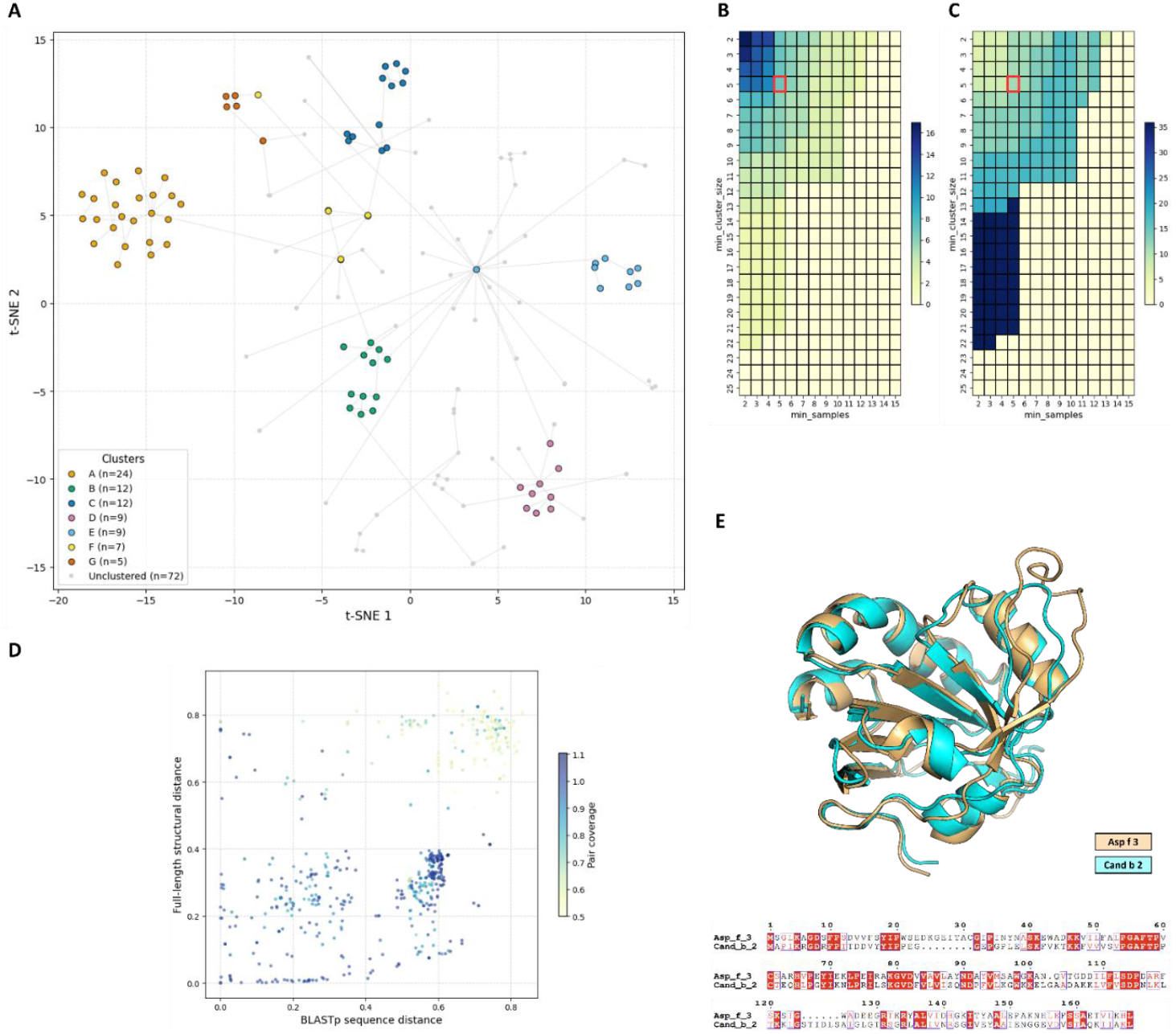
Global structure-based landscape of fungal allergens. (**A)** Global structure-based space of 150 experimentally validated fungal allergens constructed from full-length protein structures. Pairwise distances were derived from TM-scores, visualized by t-SNE, clustered using HDBSCAN, and connected by a minimum spanning tree; each node represents an allergen. **(B)** Heatmap of the number of clusters identified by HDBSCAN across combinations of *min_samples* and *min_cluster_size*, showing a progressive reduction in the number of detected clusters as parameter thresholds increase. **(C)** Heatmap of cluster size across the same combinations of *min_samples* and *min_cluster_size*, showing an initial increase in cluster size followed by collapse into unassigned points beyond specific thresholds. (D) Relationship between sequence similarity and structural similarity among fungal allergen pairs with sufficient sequence coverage (coverage > 50%). Structural distance (1 - TM-score) increased with decreasing BLAST sequence identity, showing a moderate correlation (Pearson r = 0.56; Spearman ρ = 0.62). **(E)** Representative example of structurally conserved but sequence-divergent fungal allergens. Asp f 3 (UniProt: O43099) and Cand b 2 (UniProt: P14292) exhibit high structural similarity (TM-score = 0.88) and are assigned to the same cluster (Cluster B) despite low sequence identity (33.52%) based on BLASTp analysis. Sequence divergence is illustrated by pairwise sequence alignment generated using CLUSTALW (v2.1) and visualized with ESPript (v3.2).

Local density analysis further supported the presence of structurally coherent neighborhoods within the manifold. The mean distances to the 3 and 5 nearest neighbors were 0.36 and 0.41, respectively, substantially lower than the global average pairwise distance. Consistently, the lower tail of the distance distribution showed that 25% of allergen pairs had distances below 0.7161, suggesting the existence of compact structural regions embedded within the global allergen space. To identify densely populated regions within this manifold, clustering was performed using HDBSCAN. Systematic variation of density and minimum cluster size parameters revealed that increasing either parameter led to a reduction in the number of detected clusters, while cluster sizes initially increased and subsequently collapsed into unassigned points beyond specific thresholds (Fig. 2B, C). Based on the stability of cluster number across parameter ranges, clustering was performed using a density parameter that maintained cluster structure even when the minimum sample size was increased to five.

Under these conditions, 68 out of 150 fungal allergens (45.33%) were assigned to seven distinct clusters, while the remaining allergens were not assigned to any cluster, reflecting the continuous nature of the structural space. These cluster assignments were subsequently used for epitope-level and functional analyses.

To evaluate the relationship between sequence similarity and structural similarity within the allergen manifold, pairwise comparisons were performed for allergen pairs with sufficient sequence coverage (coverage > 50%). Across these pairs, TM-score-based structural distance increased with decreasing BLAST sequence identity, showing a moderate correlation (Pearson r = 0.56; Spearman ρ = 0.62) (Fig. 2D). However, the dispersion around this trend remained substantial (RMSE = 0.21), indicating that individual protein pairs often deviated markedly from sequence-based expectations.

An illustrative example is provided by Asp f 3 (UniProt: O43099) and Cand b 2 (UniProt: P14292), which exhibited high structural similarity (TM-score = 0.88) and were assigned to the same cluster (Cluster B) within the allergen manifold, despite sharing low sequence identity (33.52%) based on BLASTp analysis (Fig. 2E). These observations highlight that structurally conserved relationships among fungal allergens can persist even in the absence of strong sequence similarity.

### 3.2. Epitope-level clustering reveals both conserved and divergent allergen families

Epitope-restricted structures were generated for all 150 fungal allergens using experimentally validated epitope prediction thresholds. On average, epitope structures retained 116.7 residues per protein, corresponding to 44.24% of the full-length allergen sequence (mean allergen length = 309.4 residues).

Using the same distance metric and clustering framework applied to full-length structures, HDBSCAN clustering of epitope-restricted structures assigned 56 allergens to four distinct epitope clusters (Fig. 3A). Despite the reduction in structural content, epitope-level clustering largely recapitulated the organization observed at the full-length protein level.

**Figure 3.**
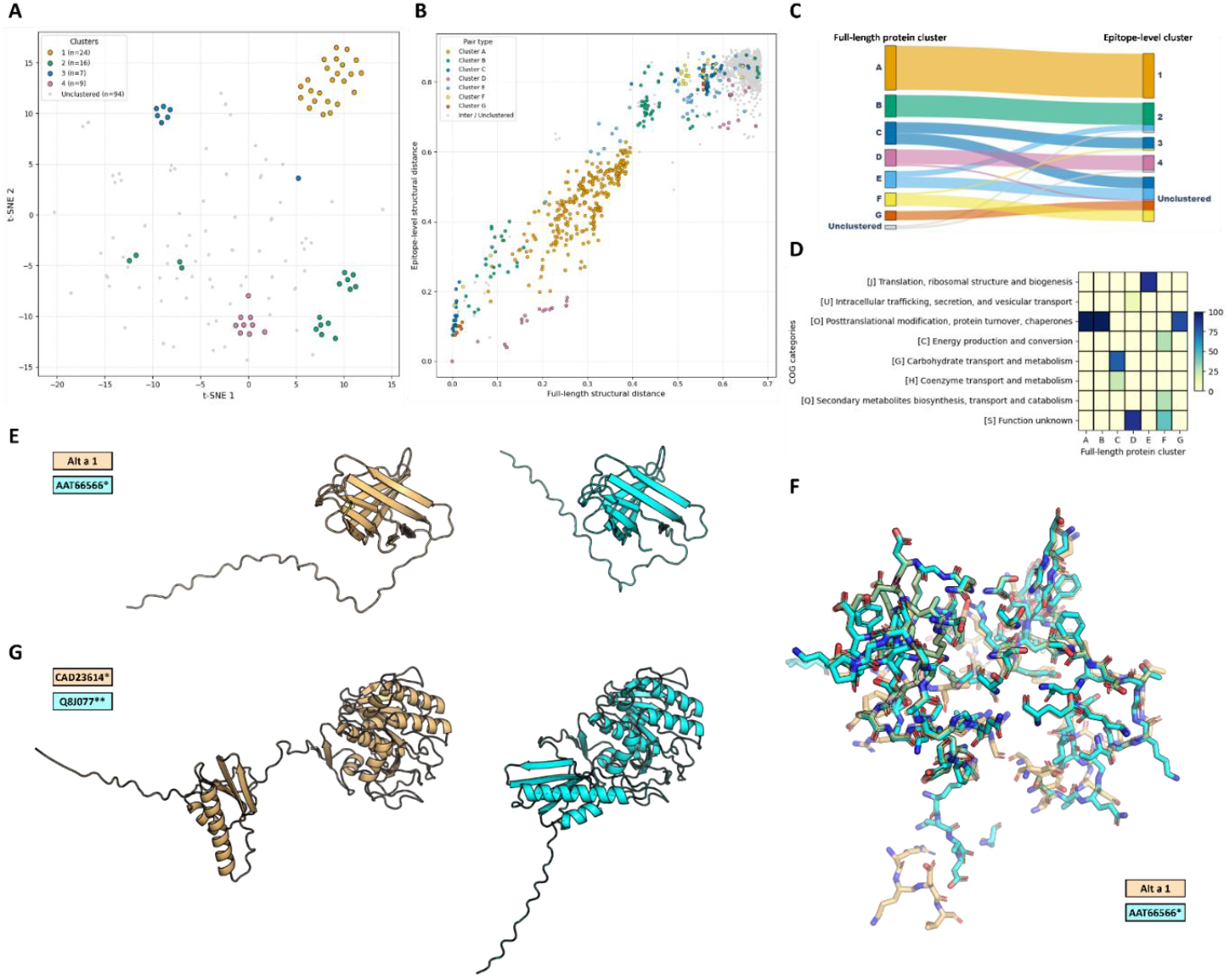
Epitope-level structural organization of fungal allergens. **(A)** Two-dimensional visualization of epitope-restricted structures derived from 150 fungal allergens. Pairwise epitope-level structural distances were computed using TM-score and visualized by t-SNE. Colors indicate epitope clusters identified by HDBSCAN, while gray points represent unclustered allergens. **(B)** Relationship between full-length structural distance and epitope-level structural distance for allergen pairs with full-length structural distances below the maximum intra-cluster threshold, showing a strong positive correlation between the two distance measures (Pearson r = 0.9521; Spearman ρ = 0.6913; RMSE = 0.0639). Colored points indicate allergen pairs within the same full-length structural cluster as defined by HDBSCAN, whereas gray points represent inter-cluster or unclustered allergen pairs. **(C)** Correspondence between full-length protein clusters and epitope-level clusters, visualized using a Sankey diagram generated with plotly (v2.29.1). Most epitope clusters predominantly originate from a single full-length cluster, while a subset of allergens is redistributed across epitope clusters. **(D)** Functional composition of full-length protein clusters based on COG category annotation derived from eggNOG-mapper. **(E)** Representative example of full-length structural similarity between Alt a 1 from *Alternaria alternata* and a structurally related protein from *Stemphylium vesicarium* (GenBank: AAT66566). **(F)** Epitope-restricted residue structures of Alt a 1 and AAT66566, shown as an overlaid licorice representation, illustrating further convergence of epitope organization relative to full-length structural similarity. **(G)** Representative example of full-length structural similarity between two subtilisin-like serine proteases from *Trichophyton benhamiae* (GenBank: CAD23614) and *Trichophyton schoenleinii* (UniProt: Q8J077.1). * and ** indicate proteins without registered allergen names in the WHO/IUIS Allergen Nomenclature database; GenBank Protein IDs or UniProt accession IDs are used instead.

Across all four epitope clusters, more than 80% of members originated from a single corresponding full-length allergen cluster, indicating substantial preservation of cluster structure at the epitope level (Fig. 3C). In particular, full-length Cluster A (n = 24) and Cluster B (n = 16) were almost entirely retained as Epitope Clusters 1 and 2, respectively. Similarly, eight out of nine members of full-length Cluster D were assigned to a single epitope cluster. In contrast, only half of the members of full-length Cluster C (n = 12) were assigned to Epitope Cluster 3, indicating reduced coherence at the epitope level for this cluster. Consistent with this observation, Cluster C also exhibited a fragmented functional distribution in COG-based analysis, in contrast to Clusters A, B, and D, which maintained coherent functional profiles under epitope-based analysis (Fig. 3D).

To quantitatively assess the relationship between structural similarity at the full-length and epitope levels, pairwise TM-score-based distances were compared for all allergen pairs with full-length TM distances below the maximum intra-cluster distance (0.68). Epitope-level structural distance showed a strong positive correlation with full-length structural distance (Pearson r = 0.9521; Spearman ρ = 0.6913; RMSE = 0.0639) (Fig. 3B), indicating that relative structural similarity among allergens is largely preserved following epitope extraction.

Representative examples further illustrated this relationship. Alt a 1 from *Alternaria alternata* and a structurally related protein from *Stemphylium vesicarium* (GenBank: AAT66566) were assigned to full-length Cluster D and Epitope Cluster 4. While the two proteins exhibited moderate full-length structural distance (0.2549), their epitope-restricted structures showed increased similarity, with a reduced epitope-level distance of 0.1830 and substantial residue overlap (Fig. 3E, F).

A similar pattern was observed for two subtilisin-like serine proteases from *Trichophyton benhamiae* (GenBank: CAD23614) and *Trichophyton schoenleinii* (UniProt: Q8J077), which were consistently assigned to full-length Cluster A and Epitope Cluster 1. Although differences in overall domain arrangement resulted in moderate full-length structural distance (0.2936), epitope-restricted structures showed closer similarity (epitope distance = 0.2701), reflecting convergence at the epitope level (Fig. 3G, Supplementary FigS. 2).

### 3.3 Manifold inclusion reveals a structurally robust cluster of allergen-related fungal proteins

To identify fungal proteins structurally related to known fungal allergens, a total of 19,999 fungal protein structures were screened against the reference fungal allergen set using MMseqs2 with relaxed sequence identity and coverage thresholds (identity and coverage > 10%). This initial screening yielded 1,352 candidate fungal proteins for downstream structural analysis.

AlphaFold-predicted structures were generated for all 1,352 candidates and subjected to manifold inclusion testing. Among these, 1,216 fungal proteins satisfied quality and inclusion criteria, and 10.36% were assigned nearest-neighbor relationships to reference allergens belonging to Cluster A (Fig. 4A).

**Figure 4.**
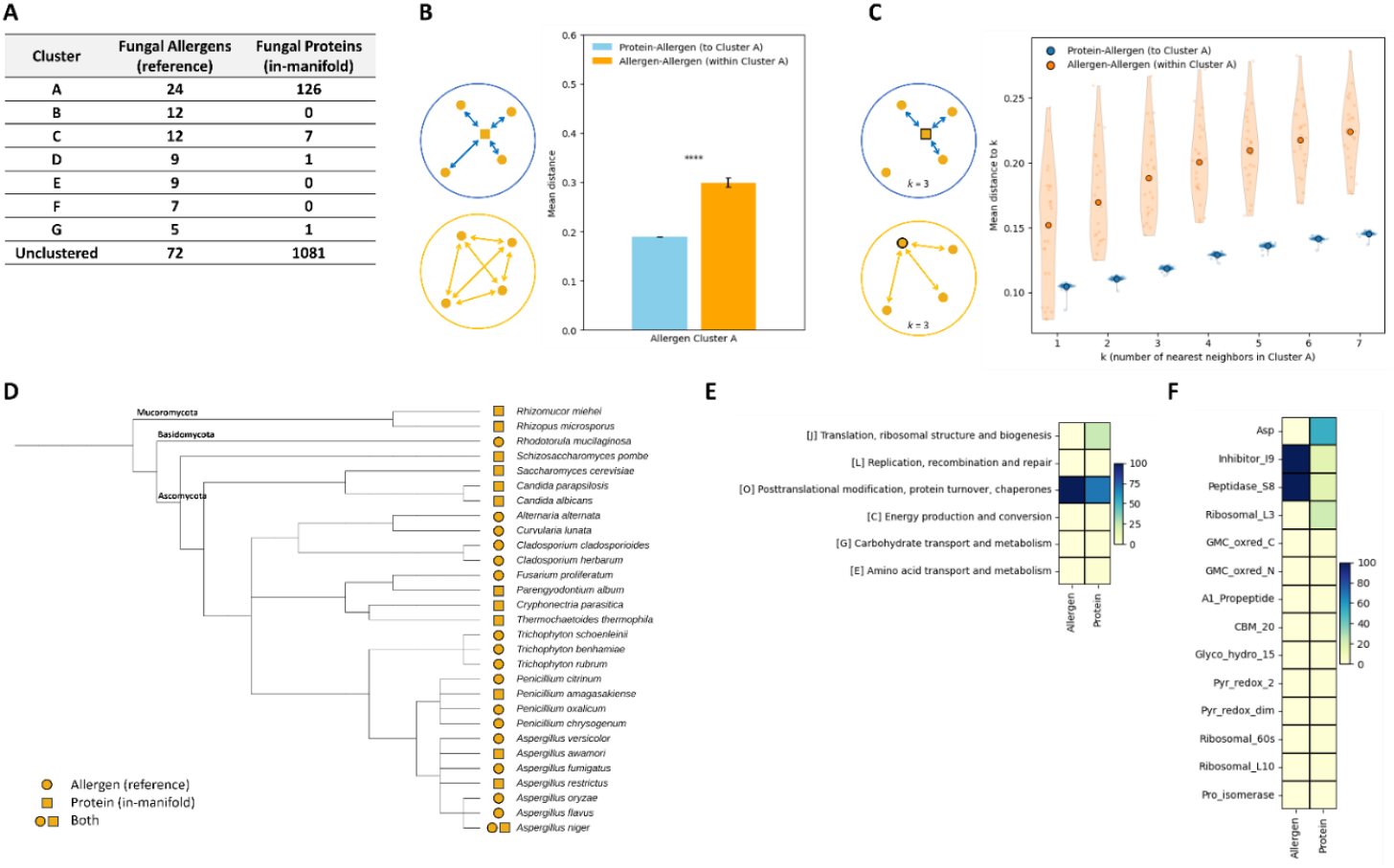
Screening and manifold inclusion of fungal proteins with enrichment in Cluster A. **(A)** Distribution of reference fungal allergens and manifold-included fungal proteins across full-length allergen clusters. A subset of fungal proteins is preferentially assigned to a specific allergen cluster, with the majority mapping to Cluster A based on nearest-neighbor manifold inclusion. **(B)** Comparison of pairwise structural distances within Cluster A. Distances between manifold-included fungal proteins and reference allergens are consistently lower than distances among reference allergens themselves, indicating tight co-localization within the same structural region. **(C)** Local structural density analysis within Cluster A based on k-nearest neighbor distances (k = 1-7). Manifold-included fungal proteins form dense neighborhoods comparable to those of reference allergens across increasing k values, demonstrating robustness of manifold inclusion. **(D)** Taxonomic distribution of species represented by reference fungal allergens and manifold-included fungal proteins. Manifold-included proteins extend the phylogenetic distribution of allergen-associated structures across multiple fungal phyla, including Ascomycota, Basidiomycota, and Mucoromycota. Circles indicate species containing reference fungal allergens, squares indicate species containing manifold-included fungal proteins, and combined symbols indicate species represented by both. **(E)** Functional composition of manifold-included fungal proteins and reference allergens in Cluster A based on COG category annotation derived from eggNOG-mapper. **(F)** Pfam domain composition of manifold-included fungal proteins and reference allergens in Cluster A, highlighting recurrent domains shared between the two groups.

Manifold-included fungal proteins assigned to Cluster A were co-localized within the same structural region as reference allergens of this cluster. Pairwise structural distances between manifold-included fungal proteins and reference allergens were consistently lower than distances observed among reference allergens themselves within Cluster A (Fig. 4B). This observation was further supported by analysis of local structural density based on k-nearest neighbor distances (k = 1-7), which showed that manifold-included fungal proteins formed dense neighborhoods comparable to those of reference allergens, independent of the chosen k value (Fig. 4C).

Functional characterization of manifold-included fungal proteins revealed a predominance of proteins assigned to COG category O (posttranslational modification, protein turnover, and chaperones; 69.05%), with a smaller fraction assigned to COG category J (translation, ribosomal structure and biogenesis; 24.60%) (Fig. 4E). Pfam domain analysis identified recurrent domains previously observed in reference fungal allergens, including Inhibitor_I9 and Peptidase_S8 (each 15.87%), as well as frequent occurrence of Asp and Ribosomal_L3 domains (Fig. 4F).

At the species level, manifold-included fungal proteins were distributed across a broader taxonomic range than species represented in current allergen databases. In addition to the 17 species previously associated with reference fungal allergens, proteins from 12 additional species were identified (Fig. 4D), spanning multiple fungal phyla including Ascomycota, Basidiomycota, and Mucoromycota.

### 3.3. Cross-kingdom allergens exhibit structural similarity relative to the fungal allergen manifold

To assess structural similarity between fungal allergens and allergens from other kingdoms, 1,289 experimentally validated non-fungal allergens were screened against fungal allergen structures using MMseqs2 with relaxed sequence identity and coverage thresholds (identity and coverage > 10%). This screening identified 72 non-fungal allergen proteins showing detectable similarity to fungal allergens.

These 72 non-fungal allergen proteins were subsequently evaluated using the same manifold inclusion criteria applied to fungal proteins. Among them, 53 proteins were classified as manifold-included (Fig. 5A). Notably, none of the manifold-included non-fungal allergens mapped to Allergen Clusters A, D, or G, suggesting that cross-kingdom structural similarity is unevenly distributed across the fungal allergen structural landscape.

**Figure 5.**
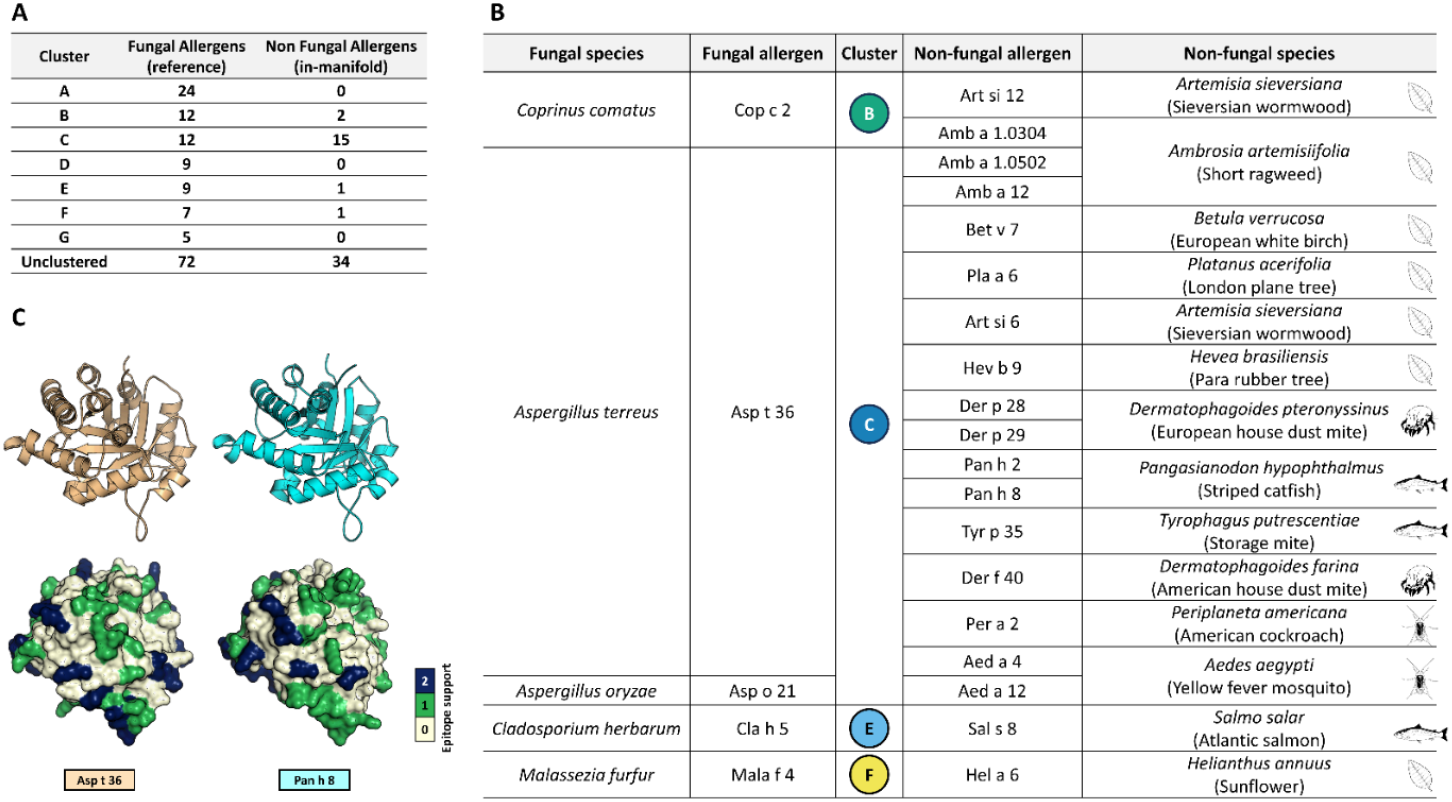
Cross-kingdom structural similarity between fungal allergens and non-fungal allergens. **(A)** Manifold inclusion analysis of non-fungal allergens. Of 1,289 experimentally validated non-fungal allergen proteins, an initial sequence-based pre-filtering using relaxed identity and coverage thresholds identified 72 candidates showing low-level sequence similarity to fungal allergens. Structural evaluation of these candidates using manifold inclusion criteria classified 53 proteins as manifold-included. **(B)** Taxonomic and allergen-level correspondence between fungal allergens and structurally similar non-fungal allergens across kingdoms. These allergens originate from diverse eukaryotic lineages, including fish, mites, insects, and plants. **(C)** Representative example of cross-kingdom structural similarity. The fungal allergen Asp t 36 (Cluster C) and the fish allergen Pan h 8 share a similar overall fold at the full-length level. Surface representations are colored according to the number of epitope prediction methods supporting each residue (BepiPred and DiscoTope), illustrating concordant epitope-level similarity across kingdoms. Icons representing non-fungal kingdoms were created with BioRender.com.

Manifold-included non-fungal allergens were derived from a broad range of eukaryotic taxa. These proteins showed structural similarity to fungal allergens originating from diverse allergenic sources, including fish, mites, insects, and multiple plant species (Fig. 5B), indicating that such cross-kingdom structural relationships are not restricted to closely related lineages.

A representative example was observed for Asp t 36, a fungal allergen from the genus *Aspergillus* assigned to Cluster C, which exhibited structural similarity to the fish allergen Pan h 8 from striped catfish (*Pangasianodon hypophthalmus*). The two proteins shared a similar overall fold at the full-length level (full-length structural distance = 0.01), and epitope-restricted structural comparison showed that high similarity was maintained at the epitope level (epitope-level structural distance = 0.19) (Fig. 5C).

## 4. Discussion

This study demonstrates the value of a structure-based framework augmented with epitope-level structural analysis for understanding the organization and diversity of fungal allergens. By constructing a global structural landscape from full-length protein structures, we show that fungal allergens are not randomly distributed but are embedded within a continuous yet heterogeneous structural space^4^. Within this space, locally dense regions correspond to well-characterized allergen families and remain highly populated when analyzed at the epitope level based on IgE-binding regions. In contrast, more diffuse regions capture gradual structural transitions and structurally related outliers. Structural distance-based clustering largely recapitulates established allergen groupings, supporting the biological relevance of a structure-centric representation that extends beyond approaches relying solely on sequence similarity.

Full-length protein-based structural clusters generally retained high local density at the epitope level, indicating that major allergen families remain structurally coherent even after restriction to IgE-relevant regions. Well-characterized allergen groups, including serine proteases, redoxin/thioredoxin proteins, and enolases, formed compact neighborhoods in the global structural manifold and continued to occupy similarly dense regions when analyzed using extracted epitope structures^18,19^. Likewise, allergens containing the Alt a 1 domain, whose molecular function remains incompletely characterized, formed a broad and stable structural cluster that was largely preserved at the epitope level^20^ (Supplementary Table 1). Consistent with these observations, pairwise structural distances computed from full-length proteins showed a strong correlation with those derived from epitope structures, supporting the view that IgE-binding features are embedded within conserved structural scaffolds despite substantial reduction of structural content^21^.

At the same time, epitope-level analysis revealed informative deviations from full-length clustering that refine this interpretation. Although epitope-based distances were generally larger than full-length distances, reflecting the reduced structural context, a subset of protein pairs exhibited increased similarity when compared at the epitope level. Such behavior is likely to arise when IgE-binding regions are concentrated within specific domains of multi-domain proteins, effectively minimizing the influence of domain arrangement that dominates full-length structural comparisons^22^. For example, serine protease allergens often contain both protease and inhibitor domains whose relative organization varies across proteins; however, IgE-interacting surfaces are predominantly localized to the protease domain, leading epitope-level comparisons to emphasize shared functional surfaces while de-emphasizing differences in global domain architecture.

In other cases, discrepancies between full-length and epitope-level clustering reflect convergence at the level of local secondary structure rather than global fold similarity^23^. A representative example is provided by the mannitol dehydrogenase allergen Alt a 8, which occupies a distinct position from enolases at the full-length structural level but becomes embedded within the enolase epitope neighborhood upon epitope-based comparison. Notably, the enolase allergen Cla h 6 emerges as one of the closest structural neighbors of Alt a 8 at the epitope level. This convergence arises despite differences in global protein architecture and is driven by the alignment of locally exposed α-helical elements that dominate the predicted IgE-interacting surfaces (Supplementary Figure 3). Additional examples of epitope-level convergence across otherwise unrelated protein folds, including cases involving ribosomal-domain-containing proteins clustering with redoxin/thioredoxin epitopes, are provided in the Supplementary Table 2.

Together, these observations illustrate how epitope-level structural analysis can reconcile apparent discrepancies between global fold divergence and immunological similarity. By selectively emphasizing IgE-interacting surfaces while attenuating the influence of peripheral or non-interacting regions, epitope-based representations capture mechanistic features of potential cross-reactivity that are difficult to explain using sequence-based or full-length structural comparisons alone.

Manifold inclusion analysis suggests that the structural definition of fungal allergens extends significantly beyond the taxonomic boundaries currently listed in allergen databases. The presence of numerous uncharacterized fungal proteins within the same structural neighborhood as experimentally verified allergens (e.g., the Peptidase cluster) supports the existence of potential annotation bias in existing allergen resources^24^. These findings indicate that allergen-like structural architectures are preserved across fungal lineages more broadly than previously recognized, underscoring the necessity of structure-based approaches in identifying potential allergen candidates.

Notably, we observed the coexistence of proteins with diverse functional annotations within the allergen manifold. This implies that allergen families are not strictly distinct functional entities but rather occupy a continuous structural space formed by shared architectures. For instance, the structural affinity of the Peptidase A1 family and ribosomal uL3-containing proteins to known allergen clusters, despite their distinct functional annotations—and despite the fact that ribosomal uL3 allergens were previously present in reference datasets but remained unassigned to specific clusters—highlights a level of structural connectivity that sequence-based homology analysis may fail to capture. These results emphasize the value of the framework in redefining the scope of potential allergens based on structural homogeneity, independent of prior experimental annotation.

Expanding the analysis beyond fungi revealed cross-kingdom structural similarities, suggesting that structural motifs associated with allergenicity may be deeply conserved evolutionarily^25^. The inclusion of certain non-fungal allergens within the fungal manifold indicates that allergenicity can be attributed to specific structural architectures that transcend taxonomic boundaries. Collectively, this structure-based framework not only reconfirms lineage-specific constraints but also elucidates widely preserved structural principles, positioning it as a conceptual model system for future pan-allergen research and novel allergen prediction.

Taken together, these results support a view of allergens as elements embedded within a continuous structural manifold rather than as members of strictly discrete classes. Building on this perspective, the proposed framework leverages epitope-level analysis to reveal gradual structural transitions and partial overlaps that are not readily captured by conventional classification schemes, while also encompassing structurally related outliers. By providing a continuous, structure-based coordinate system for allergen representation, this framework establishes a foundation for future computational and machine learning approaches that can incorporate graded structural similarity and epitope conservation, moving beyond binary allergen/non-allergen classification toward a more nuanced representation of allergen space.

## 5. Acknowledgements

This study was conducted as an independent research project and all analyses were performed on a personal computing server.

## 6. Data and code availability

The source code used for the primary analyses in this study, along with a subset of the processed data, is publicly available at GitHub (https://github.com/KyuTark/AllerScope).

## 8. Supplementary figures and tables

**Supplementary Figure 1.**
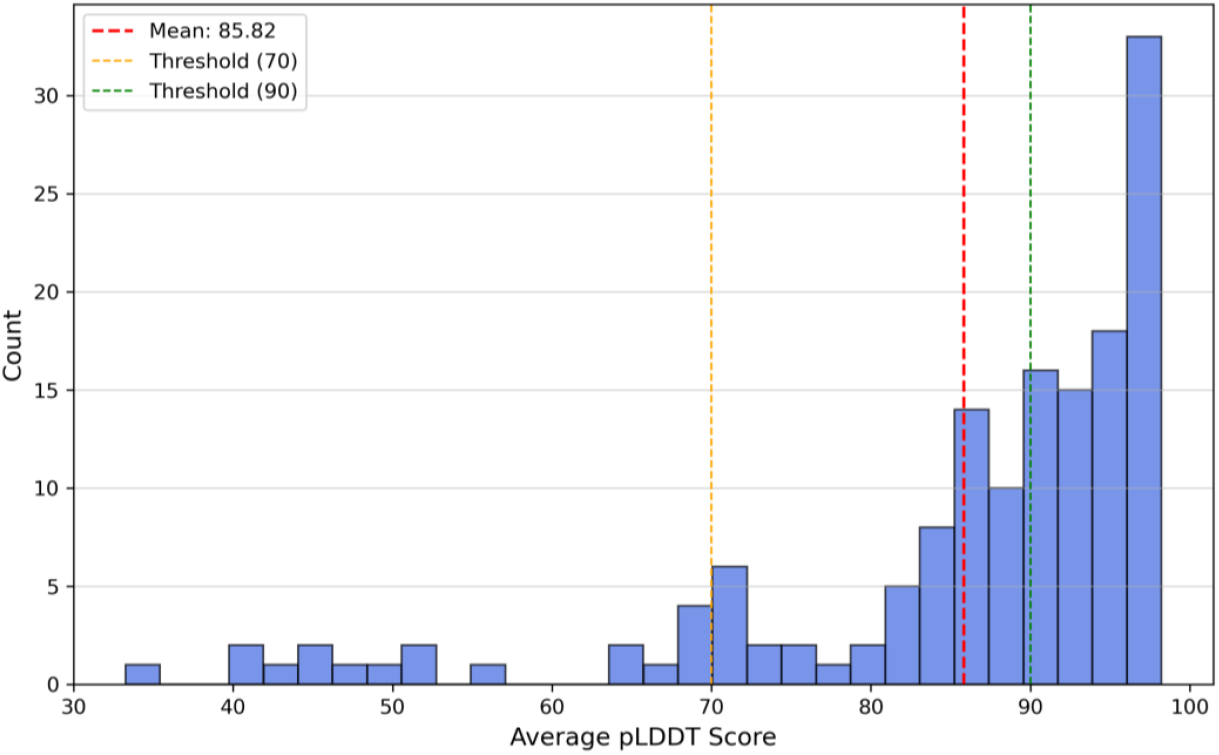
Distribution of AlphaFold predicted Local Distance Difference Test (pLDDT) scores for the 150 fungal allergen structures. The histogram displays the mean per-residue pLDDT scores for the top-ranked (rank_001) models. The global mean pLDDT score was 85.82 (red dashed line). The majority of the predicted structures (88.00%) exhibited high confidence (pLDDT > 70; orange dashed line), with 52.00% classified as very high confidence (pLDDT > 90; green dashed line). Structures with lower scores (pLDDT < 50) generally correspond to proteins containing intrinsically disordered regions (IDRs) or flexible loops, reflecting the structural heterogeneity inherent to specific allergen families rather than modeling errors.

**Supplementary Figure 2.**
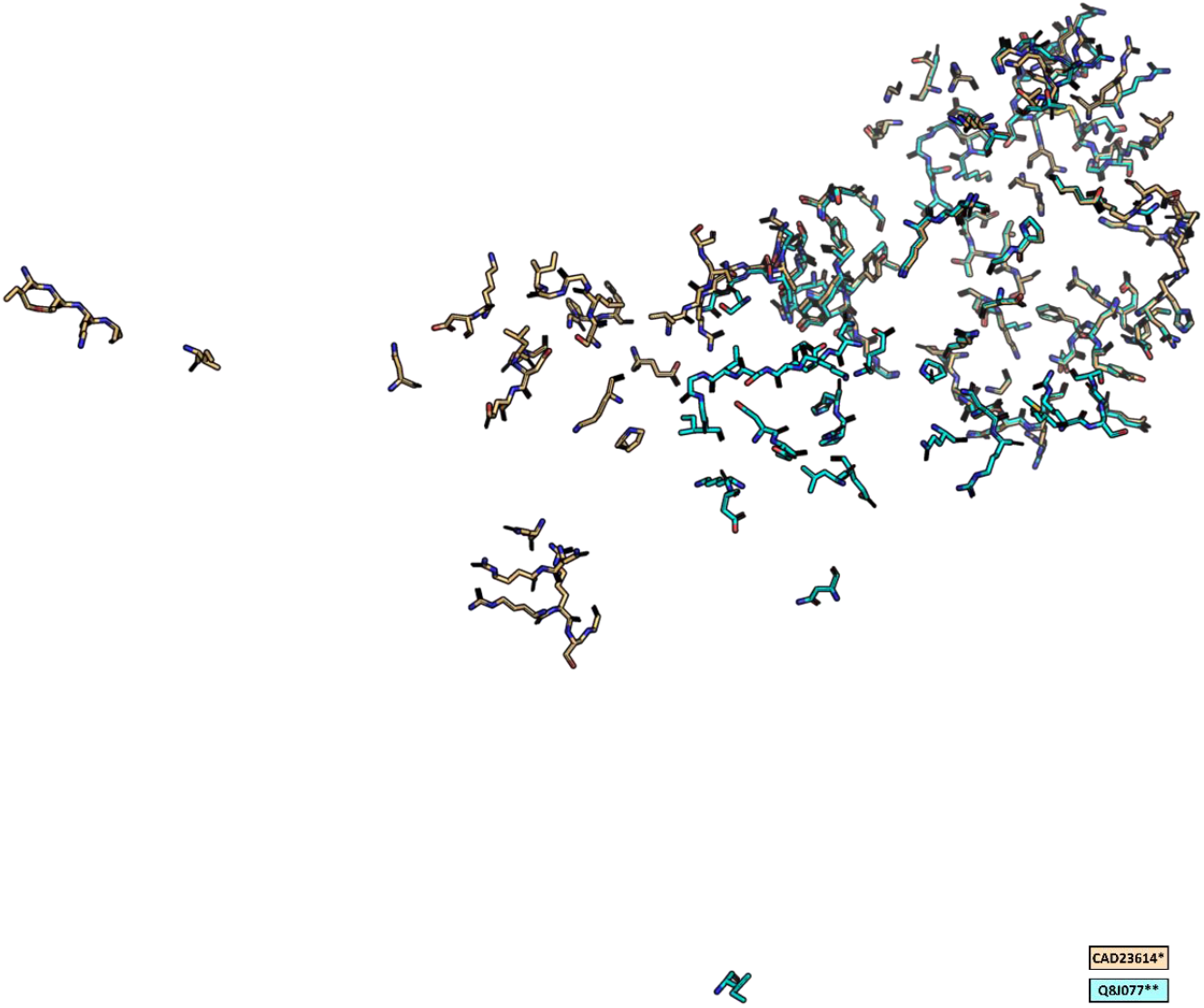
Epitope-level structural comparison of two subtilisin-like serine protease allergens from *Trichophyton benhamiae* (GenBank: CAD23614) and *Trichophyton schoenleinii* (UniProt: Q8J077). Epitope-restricted residue structures were extracted based on predicted IgE-binding regions and visualized as overlaid licorice representations. Although the two proteins exhibit moderate structural divergence at the full-length level due to differences in overall domain organization, their epitope-restricted structures show closer spatial similarity, consistent with epitope-level convergence observed within full-length Cluster A and Epitope Cluster 1. Residues corresponding to the protease domain region (upper right) were frequently detected as epitope residues in both allergens and exhibit spatial overlap, indicating localized convergence of IgE-relevant surface features.

**Supplementary Figure 3.**
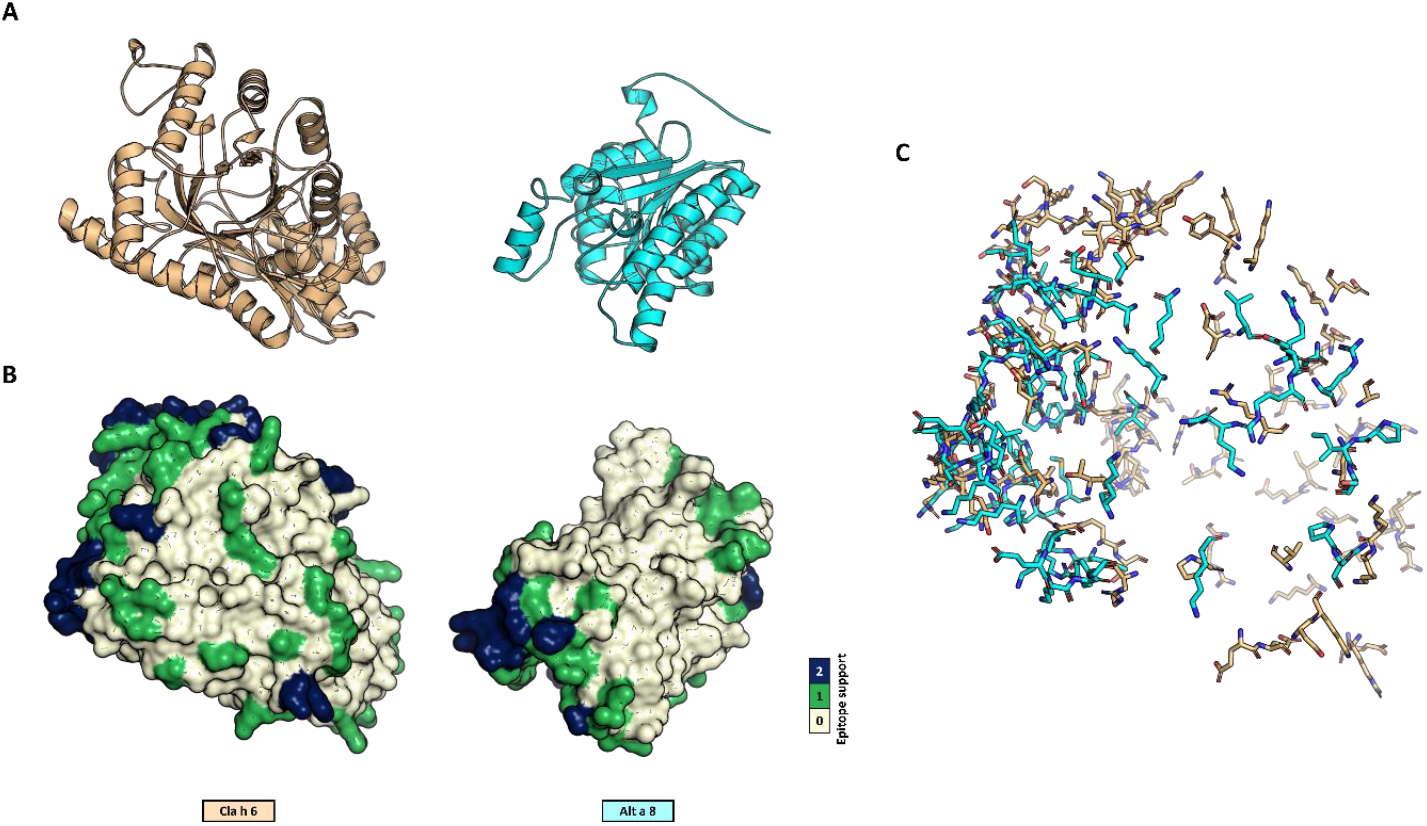
Epitope-level structural convergence between the mannitol dehydrogenase allergen Alt a 8 and the enolase allergen Cla h 6. **(A)** Full-length structural comparison illustrating distinct global folds and spatial separation of Alt a 8 and enolase proteins at the whole-protein level. **(B)** Surface representations of Alt a 8 and Cla h 6 colored according to the number of epitope prediction methods supporting each residue (BepiPred and DiscoTope), highlighting locally enriched IgE-relevant surface regions. **(C)** Epitope-restricted residue structures shown as an overlaid licorice representation, demonstrating convergence driven by locally exposed α-helical elements despite differences in global protein architecture. In contrast to full-length protein alignment, epitope-restricted structural alignment results in a shifted alignment axis that brings regions with high epitope support into a common orientation (leftward), indicating convergence guided by locally exposed IgE-interacting residues.

**Supplementary Table 1.** Pfam-based functional annotation of protein families across full-length structural clusters.

| Full-length Protein Cluster | Pfam domains |
| --- | --- |
| A | Inhibitor_I9<br>Peptidase_S8 |
| B | Redoxin<br>Thioredoxin<br>Thioredoxin_6<br>PITH |
| C | Enolase<br>TAL_FSA<br>Alpha-amylase<br>DUF1966<br>TIM<br>F_bP_aldolase |
| D | AltA1<br>NTF2 |
| E | Ribosomal_60s |
| F | adg_short_C2<br>Abhydrolase_6<br>Hydrolase_4<br>FMN_red<br>Ldh_1_C<br>Ldh_1_N |
| G | PD40<br>Peptidase_S9 |

**Supplementary Table 2.** Pfam-based functional annotation of protein families within Epitope Cluster 2, including constituent allergen proteins (UniProt ID).

| Allergen (UniProt ID) | Full-length Protein Cluster | Epitope Cluster | Pfam domains |
| --- | --- | --- | --- |
| O43099 | B | 2 | Redoxin |
| O93969 | B | 2 | Redoxin |
| P14292 | B | 2 | Redoxin |
| P56577 | B | 2 | Redoxin |
| P56578 | B | 2 | Redoxin |
| Q00002 | B | 2 | Thioredoxin, Thioredoxin_6 |
| Q1RQI9 | B | 2 | Thioredoxin |
| Q1RQJ1 | B | 2 | Thioredoxin |
| Q4WV97 | B | 2 | Thioredoxin |
| Q8TFM8 | B | 2 | PITH,Thioredoxin |
| Q9UW02 | B | 2 | Thioredoxin |
| Q9Y8B8 | B | 2 | Redoxin |
| P49148 | E | 2 | Ribosomal_60s |
| P83340 | E | 2 |  |
| Q49KL9 | E | 2 | Ribosomal_60s |
| P86254 | Unclustered | 2 | Sod_Fe_C, Sod_Fe_N |

